# Initial signs of learning: Decoding newly-learned vocabulary from neural patterns in novice sign language learners

**DOI:** 10.1101/2025.04.11.648265

**Authors:** Megan E. Hillis, David J. M. Kraemer

## Abstract

How do novice language learners represent semantic information in their new language? Whether bilinguals’ neural representations of different languages are distinct or overlapping has been well-studied, but less is known about how new knowledge is integrated into established representational networks at the earliest stages of acquisition. In addition to being an understudied case in their own right, signed languages allow the study of language learning when the sensory features of the target language are maximally dissimilar from the learners’ prior experience. Using fMRI data from 40 hearing non-signers following their first lesson in American Sign Language (ASL), we show that group-level patterns of neural representation correlate with semantic features for studied (but not unstudied) ASL signs. Furthermore, cross-language neural similarity between semantically equivalent nouns reflects individual-level comprehension of ASL. Item-level comprehension was decodable in regions that represent semantic relationships within ASL, suggesting that representations of newly acquired ASL nouns become more similar to well-known English counterparts as a result of early-stage learning. These results demonstrate the role of distributed neural regions, including perisylvian language areas as well as right parietal areas, in representing semantic content across language and modality in novice learners.

**Research Transparency Statement:** The authors declare no conflicts of interest. This research was supported by National Science Foundation Award #1822819. No artificial intelligence assisted technologies were used in the creation of this article. This research received approval from the Dartmouth Committee for the Protection of Human Subjects. No aspects of the studies were preregistered. Some stimulus videos cannot be shared due to copyright restrictions but can be viewed on the original website (www.asl-lex.org). Deidentified primary data and analysis scripts are available upon request to the corresponding author.

## Introduction

How do novice learners represent semantic information in their new language? The extent to Whether bilinguals’ neural representations of different languages are distinct or overlapping has been the topic of much research, but less is known about how new knowledge is integrated into established representational networks, especially in adults at the earliest stages of learning. Furthermore, most investigations have compared languages which share a modality (i.e., spoken languages). Spoken language users learning a signed language are an interesting case where the sensory features of the new language are maximally different from the learners’ prior experience. Interest in sign language education is on the rise in many countries including the United States^1^. Yet, despite the growing demand and the documented importance of sign language access^2,3^, very little research has explored the neural underpinnings of the early stages of sign language acquisition.

Research in bilinguals has found evidence for both commonality and divergence between neural processes underlying multiple languages in the brain, and it appears that for non-native languages these complex interactions are sensitive to age of acquisition and proficiency: a series of meta-analyses have demonstrated that in bilinguals, separate languages rely on similar networks in the brain but may utilize them in subtly different ways.^4,5^ Multivariate pattern analysis methods such as Representational Similarity Analysis (RSA)^6^ have provided a way to probe neural activity patterns for this dissociation by measuring the correspondence of fine-grained patterns of activity with an a priori model of patterns of semantic features. Semantic representations should be sensitive to conceptual properties of stimuli (e.g. object category) regardless of the modality of the stimulus cue. It is well-established that such methods can decode conceptual representations that are shared between words and pictures^7–10^ as well as written and audio presentations of the same content^11–14^. This property also extends to homologous words across languages. Cross-language decoding techniques have revealed language-independent representations of fine-grained distinctions between nouns especially in left temporal regions^15,16^, that certain tasks can induce cross-language decodability in sensory cortices (e.g. occipital regions for word reading, pre-and postcentral regions for speech production^17^, and that cross-language representations may be more strongly evoked by deeper semantic processing^18^.

These studies demonstrate that bilinguals access semantic meaning in a way that is at least partially language-independent. However, less is known about how these partially overlapping representations develop as individuals learn a new language. Some studies have found that late and less proficient bilinguals exhibit more overlap across languages than native bilinguals^19–21^, which could be attributed to use of established native language networks to scaffold new learning.

Only one study to our knowledge has utilized this type of cross-decoding in *bimodal* bilinguals – those fluent in at least two languages that differ in modality (speech versus sign). Evans and colleagues^22^ presented bilinguals of spoken English and British Sign Language (BSL) with clips of both languages during functional Magnetic Resonance Imaging (fMRI) scanning. Using RSA^23^, they probed for neural activation patterns reflecting both item-level semantic relationships and broader category-level relationships between the stimuli in each language. They found evidence of modality-specific semantic representation for each language at the item level, but only representations of broad categories (e.g. animals vs. fruits vs. vehicles) across languages. The former is consistent with a large body of research suggesting that, overwhelmingly, neural mechanisms underlying linguistic processing of sign and speech are very similar (for review, see MacSweeney and colleagues^24^), and that left inferior frontal^25–30^ and superior temporal^31–35^ areas serve linguistic functions regardless of language modality. However, given that cross-language representations did not correlate with an individual item model, the authors propose that in native bimodal bilinguals, representations of individual nouns are only shared between sign and speech at the broader conceptual level. Instead, fine-grained item information may be represented in a way that is language-specific. This is convergent with robust univariate evidence for both shared and language-specific neural underpinnings of sign comprehension^35^.

However, it remains untested how cross-language representations develop in late and novice signers. Compared to early signers, hearing late signers make less use of right parietal areas implicated in spatial processing of sign^36–38^, and show neural patterns that overlap with those evoked by their native spoken language^21,39^ similarly to novice learners of spoken languages who may use their established native language as a scaffold for subsequent learning^19,40^. Neural overlap between sign and speech also appears to increase with proficiency in the early stages of sign language learning. For example, a univariate fMRI study by Williams, Darcy, and Newman^41^ found that in participants scanned repeatedly over the course of 10 months of American Sign Language (ASL) training, lexico-semantic processing of ASL in the left inferior frontal gyrus (IFG), supramarginal gyrus (SMG), and putamen increased at each timepoint.

However, the inferences that can be drawn from similarities in univariate activation are limited. Multivariate pattern analysis techniques such as RSA, whereby the relationships between stimuli are modeled in a high-dimensional space and the presence of neural activity with a similar pattern of relationships is assessed, can provide uniquely fine-grained insights into the content of linguistic processing^20^. We leverage methods pioneered in studies of concept knowledge^42,43^, especially in physics and engineering^44–47^ to evaluate the correspondence between individual-level neural “scores” and learning measured by traditional pencil-and-paper tests.

In the present study, hearing English speakers completed a brief (∼45 minutes over two sessions) lesson in ASL, then participated in fMRI scanning during which neural activity was recorded while they watched clips of concrete nouns that they had seen during the lesson or new, unstudied signs. They also watched audiovisual clips of translationally equivalent nouns in English. At three time points (after training on days 1 and 2, as well as an additional quiz administered three weeks later), they completed a free recall task, translating each sign into English. An overview of the training procedure as well as recall performance for the studied items is illustrated in Figure 1. For ASL and English separately, we identified a network of neural parcels which were sensitive to within-language semantic relationships, using a method similar to the broad conceptual model utilized by Evans and colleagues^22^. If later and less proficient learners activate their established language network while processing their target language, we might expect that the fine-grained item information that was found to be language-specific in bimodal bilinguals might be more shared in novices. Furthermore, if these shared representations are a meaningful indicator of learning, we hypothesize that the degree of overlap should vary with individual-level comprehension of semantic information from ASL.

**Figure 1.**
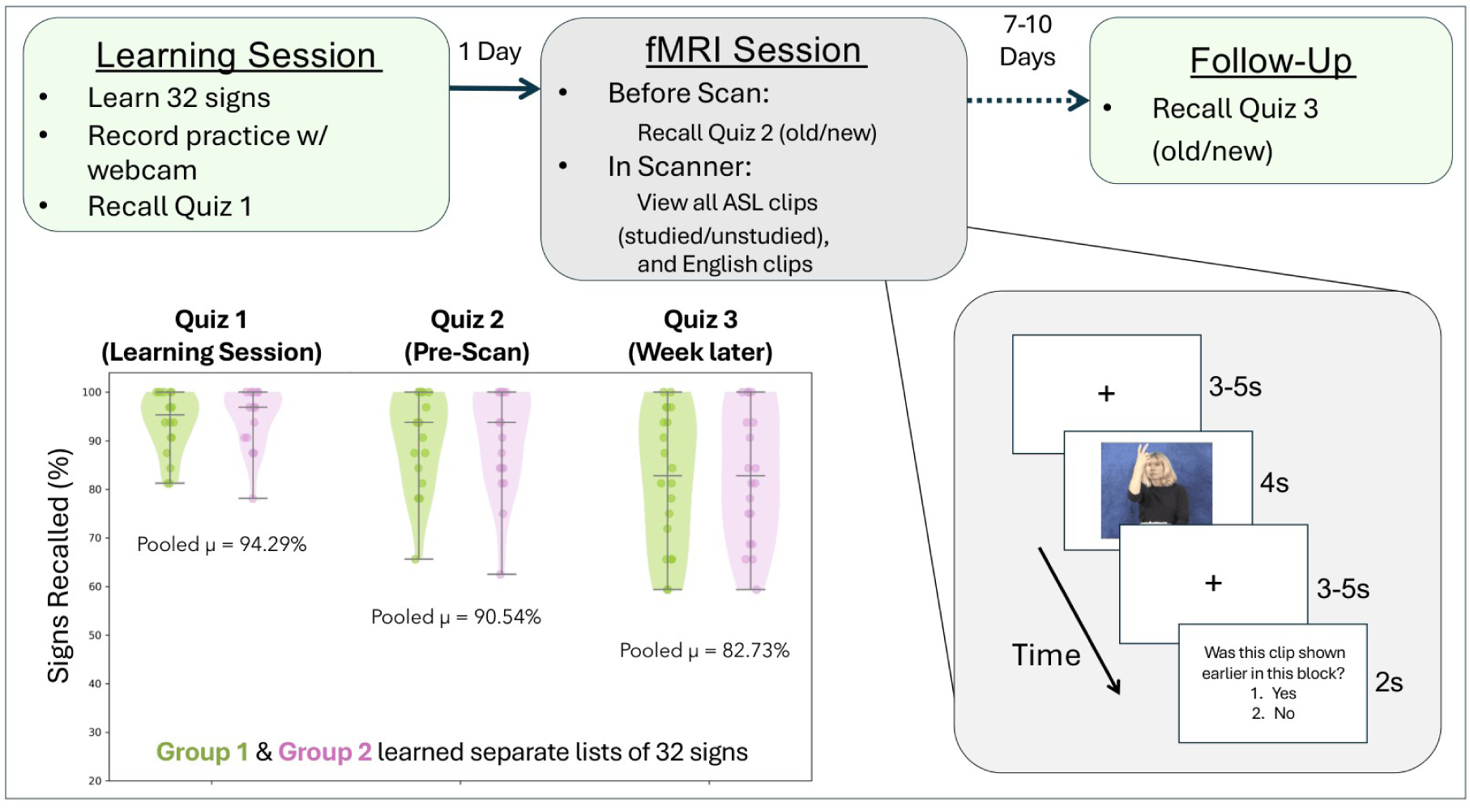
Learning procedure and recall quiz scores. Note. ASL learning procedure and behavioral recall scores. Participants completed an online lesson one day prior to the scan session, which concluded with a free recall quiz. One day later, they completed a brief review and a second quiz immediately before fMRI scanning, where they were shown single noun clips in ASL and English, sometimes followed by an attention check. Finally, one week after scanning participants completed a third recall quiz. Individual quiz scores are shown as a scatter plot overlaid on the violins. The horizontal lines indicate the maximum, mean, and minimum score at each timepoint, while the curves represent the frequency of the result in the distribution. Screenshots of the ASL stimulus videos shown in the fMRI procedure schematic are credited to the ASL-LEX project (https://asl-lex.org/), used with permission.

## Methods

### Participants

Forty-three volunteers including both undergraduate students and members of the broader community participated in this study. All participants were fluent English speakers with no prior training in any signed language. Three were excluded (one due to failing to complete the pre-scan quiz, and two due to excessive motion in the scanner) for a final N = 40 (28 female, 2 nonbinary, mean age=20.3 years, SD=3.0). Participants provided informed consent at the beginning of each session and were compensated either with curricular extra credit points or a small monetary reward. All protocols were approved by the Dartmouth Committee for the Protection of Human Subjects.

### Stimuli and Design

Stimuli for the behavioral and scanner tasks were short video clips each containing a single noun in ASL or English. The ASL videos were provided by ASL-LEX, a database of lexical and phonological properties of ASL signs^48,49^. English videos were created by lab volunteers with efforts to mimic the style of the ASL-LEX videos. Each clip consisted of the presenter seated before a neutral background, demonstrating a single noun towards the camera. To ensure that participants would not be able to guess the meanings of the clips without training, ASL signs which were rated greater than average in transparency (the extent to which a sign’s meaning is knowable by non-signers due to iconicity, or form that resembles meaning) by a sample of hearing non-signers collected by the creators of the ASL-LEX corpus were excluded. Additionally, all ASL stimuli were pilot tested on Amazon Mechanical Turk. A group of 15 English-speaking non-signers were shown each clip and asked to guess what they thought the sign could mean in English. Any signs for which any pilot participant guessed correctly were also excluded from the stimulus set.

Participants were divided into two groups of ASL learners who were each assigned to learn a different vocabulary list. Thus, for each group, there was a set of 32 studied signs, and a separate set of 32 signs which were never shown during the training phase. In each list, 12 out of the 32 signs were members of one of the three target categories (animals, fruits, and vehicles).

### Procedure

Participants completed an online vocabulary lesson (approx. 45 minutes) on the day preceding the fMRI scan. The lesson was administered through Qualtrics (Qualtrics, Provo, UT) and consisted of four blocks of eight ASL signs. Two counterbalanced groups saw different lists of 32 signs. During each block, participants watched each sign clip a few times (first slowed to half speed, then full speed) and were encouraged to mimic the video themselves. They then completed a set of multiple-choice questions and used their devices’ camera to record and upload videos of themselves practicing each sign. At the end of the four blocks, they completed a final recall task where they were asked to type in an English translation for each of the 32 items, which made up their Quiz 1 score. Copies of the lesson and quiz materials (excluding the videos which are subject to copyright) are available on Open Science Framework (https://osf.io/4bh5p).

The following day, participants completed the fMRI session. Prior to entering the scanner, they took another brief recall quiz. In this version, participants were shown all 64 signs in an old-new paradigm, where they were asked to indicate whether they had studied each sign, and for each sign marked “old”, they were asked to type in the English translation. The proportion of studied signs for which they provided the correct translation was calculated as their pre-scan (Quiz 2) recall score. In the scanner, after the initial anatomical scan, twelve clips from each of the studied and unstudied lists were presented in a randomized order in three functional runs. Each clip was shown twice per run, resulting in six presentations each. On a small number of trials (10%), the video clip was followed by an attention probe (“Have you seen this clip already during this run?”) which participants answered by pressing a button with their right index or middle finger. After three functional runs of ASL clips, participants repeated this same task in English. English was presented at the very end to avoid informing participants of the full set of nouns, which might help them guess the meaning of unstudied signs.

One week after the fMRI session, participants were contacted by email and asked to take the same old-new recall test a final time (Quiz 3) to assess longer-term retention of the signs. The complete timeline of the three time points and quiz performance at each are diagrammed in Figure 1.

### fMRI Data Acquisition and Preprocessing

Brain images were acquired using a 3 Tesla Siemens PRISMA fMRI scanner with a 32-channel head coil. A single high-resolution T1-weighted anatomical scan and six functional runs of 241 measurements (6.3 minutes) were performed for each participant with a 240 mm2 field of view to provide full brain coverage over 46 slices (Flip angle = 79°; TE = 32 ms; TR = 2500 ms; 3x3mm voxels). In the scanner, stimuli were presented using PsychoPy^50^ version 2021.2.3 (using Python 3.6).

#### Image Preprocessing and General Linear Model

Brain images from both datasets were preprocessed with fMRIprep version 21.0.1^51,52^, which is based on *Nipype* 1.6.1^53^. As recommended by the software developers, boilerplate text generated by fMRIprep is reproduced exactly in the Supplementary Materials. In summary, preprocessing included skull stripping, slice-time correction, smoothing, registration and normalization. fMRIprep also estimates the time course of confounds such as head motion, which are discussed below.

##### Beta Estimation

After preprocessing, a univariate regression model using the GLM was then calculated at the trial level, such that beta-value estimates for each stimulus and their temporal derivatives were generated separately for each run using Nilearn’s FirstLevelModel tool^54^. Additional regressors of no interest were also included in the model: the time course of six motion parameters (translation and rotation in the X, Y, and Z dimensions) generated by fMRIprep, and spike regressors for any time points with framewise displacement greater than three standard deviations above the mean. Finally, for stimuli which appeared in more than one run, beta estimates were additionally combined across runs with an item-level fixed effects model, yielding a single contrast estimate for each item in each language.

##### Dissimilarity Matrices

To assess the correspondence between neural activity patterns and the hypothesized relationships between stimuli, we constructed neural dissimilarity matrices (DMs) of the correlation distances between the neural responses to each item in each language. One DM was constructed for each subject at each parcel in the Schaefer cortical parcellation atlas^55^ (500-parcel version).

### Pooled RSA for Parcel Selection

As a region selection step, we first sought to identify a subset of brain regions which were generally sensitive to semantic relationships between the items across the group. However, we expect that individual differences in ASL comprehension (the exact variation in individual learning that this paradigm was designed to test) could add significant noise to the group-level signal. Thus, at this region selection step, we pooled these participants’ data with a sample from previous work (N=10)^56^. These participants were also hearing non-signers who completed a nearly identical learning paradigm with stimuli drawn from the same superset, but with more training time such that the majority of participants learned all signs to ceiling. By including data from novice learners who understood the ASL content well, we hope to select brain regions where we can reasonably expect successful learning to give rise to decodable semantic information in the relationships between the ASL items.

#### Semantic and Perceptual Feature Models

##### Word2Vec

To model semantic relationships between the stimuli, we calculated the representational distance between each pair of items using a word2vec model^57^. Cosine dissimilarity between each pair of target items was calculated with pretrained embeddings derived from the full corpus of English wikipedia^58^. Particularly given that these participants were non-signing English speakers, we expect the resulting dissimilarity matrices to capture meaningful semantic relationships between the items regardless of language.

##### Sign Form Features in ASL

As with most languages, phonology in ASL is not necessarily orthogonal to semantics. We approach this potential confound in two ways: first by excluding highly transparent signs (as described under ‘Stimuli’), and secondly by modeling lower-level sign form features using metadata codes provided by the ASL-LEX 2.0 database^49^. These codes were created based on the Prosodic Model^59^ to describe the structural composition of signs, encompassing handshape, palm orientation, location in sign space, movement, and sign type (one- or two-handed, symmetry) as well as other features. We calculated sign form similarity between two signs as the proportion of these features that each pair of signs shared.

##### Auditory Features in English

We modeled phonological similarity between the English stimuli with a similar procedure. The phonetic algorithm Metaphone^60^ was used to calculate the Levenstein distance between each pair of English words.

The dissimilarities between the stimuli estimated by each model were first z-scored and rescaled from zero to one, where zero indicates the exact same item and one indicates the maximal distance. Then prior to RSA the relevant models for each language were orthogonalized to each other using linear regression, as diagrammed in Supplementary Figure 1A.

After computing the item-level neural dissimilarities in each parcel as described above, we used RSA to assess the correlation between the neural data and the word2vec model (after orthogonalization to sign form features). Spearman correlation between the parcel-level DM and the model was passed through a Fisher z-transformation and compared to a null distribution calculated as the dot product of the true model and a randomly permuted model standardized over 1,000 iterations. The resulting correlations were subjected to a one-sample T-test at each parcel to identify brain regions where the neural correlation with the semantic model was greater than zero, in other words, areas where BOLD activity patterns reflected semantic relationships between the items. Finally, the T-test results for each of the 500 parcels were submitted to a sign-flipped permutation test, by which each observed T-statistic is repeatedly compared to a distribution of scores from other parcels which have been randomly sign-flipped^61^ to control for multiple comparisons. Parcels with a corrected p<0.05 are shown in Figure 2A, and were used as a binary mask for all further analyses. Heatmaps of all parcels which met these criteria are shown in Supplementary Figure 1B (for ASL trials) and 1C (for English trials).

**Figure 2.**
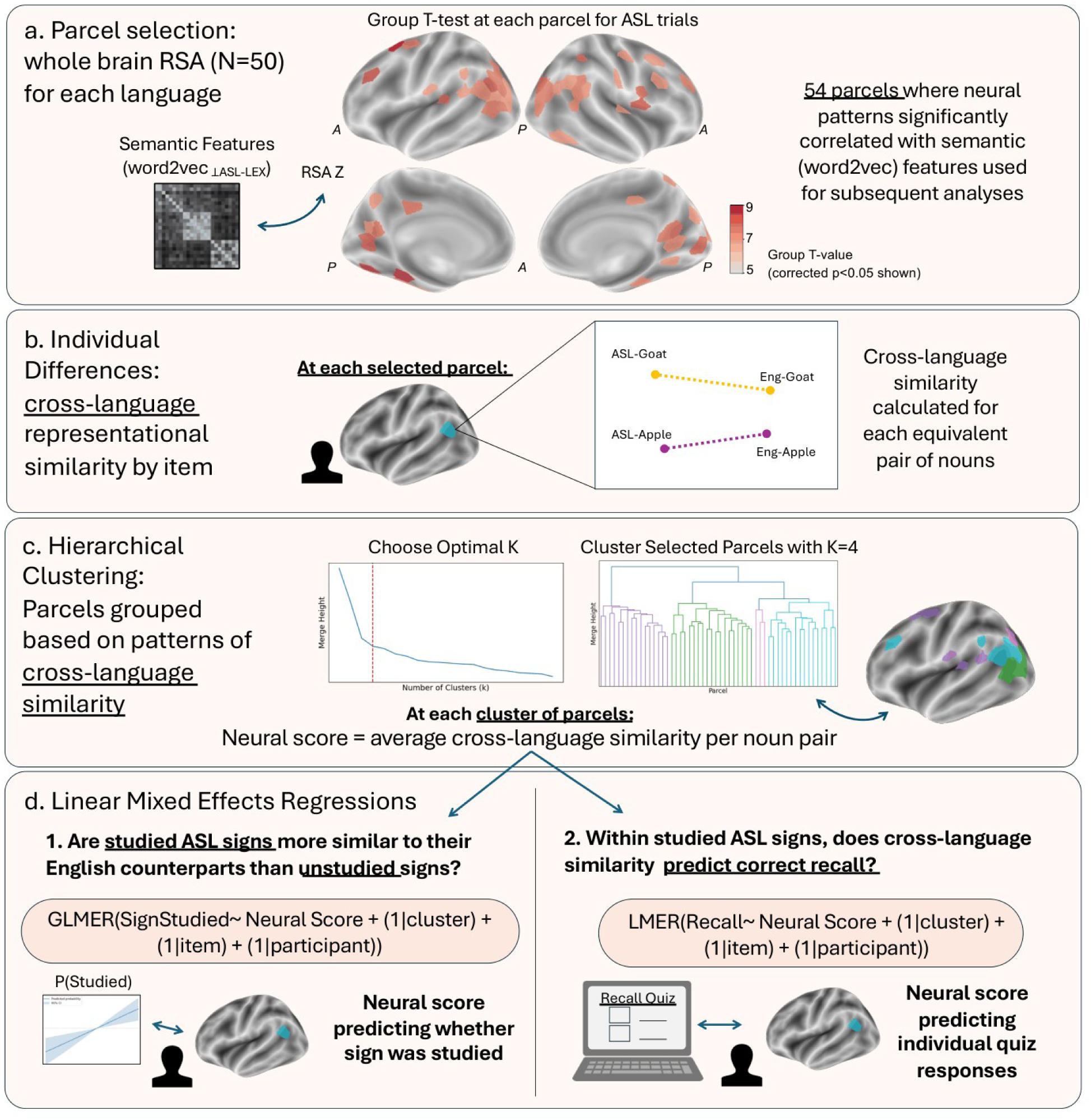
Analysis procedure. Note. Analysis procedure for ASL trial RSA mask. The same procedure was then repeated for the significant parcels from the English RSA. A. Group-level correlation of ASL neural data to semantic model (which had been orthogonalized to phonological features) was assessed with RSA. Parcels for which pooled (N=50) RSA group T-statistics were significant (corrected p<0.05) were used for subsequent analyses. B. In each of these parcels, we calculated the cross-language representational similarity between each equivalent pair of nouns for each participant. C. These item and participant-level similarity scores were used to hierarchically cluster the 54 selected parcels, yielding a map of brain regions that respond similarly to patterns across the stimulus set. We assessed the optimal dendrogram cut point using the acceleration method (using the second derivative of change in merge height to measure the elbow of the curve). The optimal solution yielded four clusters. The average score for each item and participant in each of these clusters is used as a “neural score”. D. We then used linear mixed effects models to assess whether these networks meaningfully differentiated between studied and unstudied signs, and whether they were sensitive to individual recall performance for studied signs.

### Hierarchical Clustering

The RSA parcel selection described above yielded 54 significant parcels for the ASL trials and 24 significant parcels for English. Because this is a still relatively high number of features relative to observations (12 target items per subject in each language), we next sought to group those parcels into clusters using hierarchical clustering. We computed the optimal number of clusters (K) for each solution by taking the second derivative of the change in merge height for each value of K (known as acceleration)^62^, and used this to confirm the elbow of this plot, which resulted in four clusters in both cases. Lists of the precise parcel numbers which made up each cluster are reported in Supplementary Table 1. Crucially, while the parcels were originally selected using only within-language neural DMs, this step creates clusters of regions with similar across-language patterns of neural similarity. We computed an individual participant “neural score” for each item and cluster by averaging the cross-language similarities for each parcel in each cluster.

## Results

### Behavioral Performance

Both groups learned the studied signs with fairly high accuracy. Figure 1 shows the target item recall scores for the two groups for all three timepoints (M_T1_=94.3%, SD_T1_=1.0%; M_T2_=90.5%, SD_T2_=1.6%; M_T3_=82.7%, SD_T3_=2.1%). A two-way mixed ANOVA confirmed that the two groups performed comparably at each time point (F_group_=0.15_(38,76),_ p=0.70), therefore we analyze them as a single, pooled sample. For each participant and item, recall score was calculated as the average of the three quizzes.

### Cross-language similarity in ASL-sensitive regions

After computing neural scores for each participant, item, and cluster as described above, we assessed the relationship between these scores and behavioral measures in two ways. First, we fit a logistic regression model to predict whether a given sign was among the subset of signs that each participant studied during the lesson phase. Then, we used a gaussian linear mixed effects model to estimate the relationship between the neural similarity of studied signs to their English counterparts and individual recall. The latter is the most direct measure of individual differences: it indicates the extent to which ASL signs with greater neural similarity to their English counterparts were more likely to be recalled while controlling for participant, item, and cluster effects. Both of these analyses were implemented using Pymer4^63^, a Python implementation of the R package lme4^64^. Pymer4 provides Satterthwaite confidence intervals and significance values which are computed using a bootstrapped estimation of degrees of freedom, helping overcome traditional limitations of interpreting linear mixed effects^65^.

#### Neural score predicting whether sign was studied

A logistic (binomial) linear mixed effects regression model predicting whether each sign belonged to that participant’s studied sign list was fit with neural score as a fixed effect and participant, item, and cluster included as random effects with random intercepts. The resulting beta value of neural score predicting whether the sign was studied was significant across the entire network (β=0.18, p=0.001), and output of the model including confidence intervals is plotted in figure 3B.

**Figure 3.**
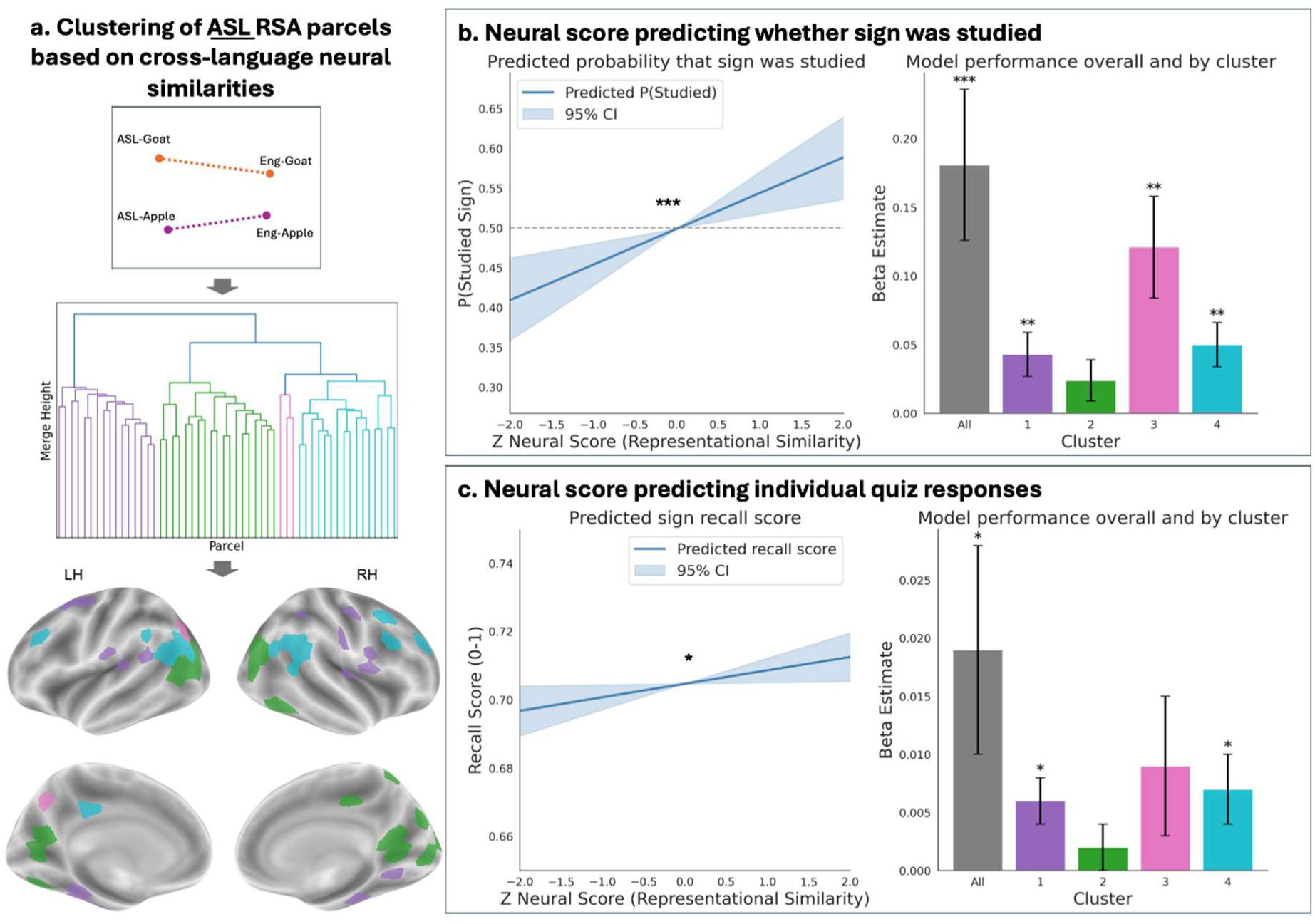
Cross-language similarity predicting individual differences in ASL RSA regions. Note. Linear mixed effect model results for ASL RSA regions. A. Diagram of hierarchical clustering procedure. A color-coded map of the parcels belonging to each of the four clusters is displayed on the semi-inflated FSAverage6 surface. B. Neural score predicting whether each sign belonged to the participant’s studied sign list. The change in predicted probability that the sign was studied over the range of neural scores is plotted on the left, while individual beta estimates of neural score for the entire model as well as split by cluster appear on the right. C. The same results are presented for neural score predicting individual recall quiz score. Bars represent standard error and are marked with asterisks by significance (p<0.05=*, p<0.01=**, p<0.001=***).

As a post-hoc analysis, we also sought to measure how the individual clusters were contributing to this effect. We fit another, identical model within each cluster, including a neural score for each parcel within the cluster and including random intercepts for participant, item, and parcel. Marginal R^2^ for each model was computed following Nakagawa and colleagues^66^ and corrected with a False Discovery Rate threshold (Benjamini-Hochberg procedure). The results are plotted in Figure 3B and summarized in Table 1.

**Table 1.** Linear Mixed Model Results for ASL RSA Regions.

| Model | $\beta$ | CI | Marginal $R^2$ | p-adj |
| --- | --- | --- | --- | --- |
| **Predicting studied sign** |  |  |  |  |
| All ASL Clusters | 0.18 | 0.07-0.29 | 0.003 | **0.001\*\*\*** |
| ASL Cluster 1 | 0.04 | 0.01-0.08 | 0.001 | **0.009\*\*** |
| ASL Cluster 2 | 0.02 | -0.01-0.05 | >0.001 | 0.104 |
| ASL Cluster 3 | 0.12 | 0.05-0.2 | 0.004 | **0.004\*\*** |
| ASL Cluster 4 | 0.05 | 0.02-0.08 | 0.001 | **0.004\*\*** |
| **Predicting recall score** |  |  |  |  |
| All ASL Clusters | 0.019 | 0.00-0.04 | 0.002 | **0.033\*** |
| ASL Cluster 1 | 0.006 | 0.00-0.01 | 0.001 | **0.024\*** |
| ASL Cluster 2 | 0.002 | 0.00-0.01 | >0.001 | 0.415 |
| ASL Cluster 3 | 0.009 | 0.00-0.02 | 0.001 | 0.199 |
| ASL Cluster 4 | 0.007 | 0.00-0.01 | 0.001 | **0.024\*** |

*Note. Normalized beta estimates, confidence intervals, marginal $R^2$ , and adjusted p-values are provided for the overall model as well as each within-cluster model for the regions selected with*
the ASL RSA. Confidence intervals are computed with Satterthwaite degrees of freedom estimation, and these are used to estimate the $p$ -values. When relevant, $p$ -values are additionally corrected with an FDR threshold. It should be noted that the prediction of studied from unstudied signs is a binomial (logistic) model, while recall score is predicted with a gaussian model, so effect sizes are not directly comparable between the models.

#### Neural score predicting recall of studied signs

Although it is interesting that studied and unstudied signs are dissociable broadly in the neural data, we also sought to test the effect of individual-level learning outcomes on this neural score. Using only data from studied signs (thus, signs that participants had seen before and had been given an English translation for but varied in how accurately they remembered that translation), we fit a linear mixed effects model to predict recall score from neural score, again including participant, item, and cluster as random effects. The result was again significant (β=0.19, p=0.033). As before, we then tested the influence of each of the four clusters with an FDR-corrected (Benjamini-Hochberg procedure) series of cluster-level mixed effects models. The results are summarized in Figure 3C as well as Table 1.

### Cross-language similarity in English-sensitive regions

The same procedure was then repeated with the network of parcels 24 identified as sensitive to semantic relationships between the items in the English trials (colored by cluster in Figure 4A, as well as plotted as a heatmap of RSA scores in Supplemental Figure 1C). After computing neural scores for each participant, item, and cluster, we fit the same two types of models: a logistic (binomial) regression to predict whether a given sign was studied, and a linear regression to estimate the relationship between the neural similarity of studied signs to their English counterparts and individual recall.

**Figure 4.**
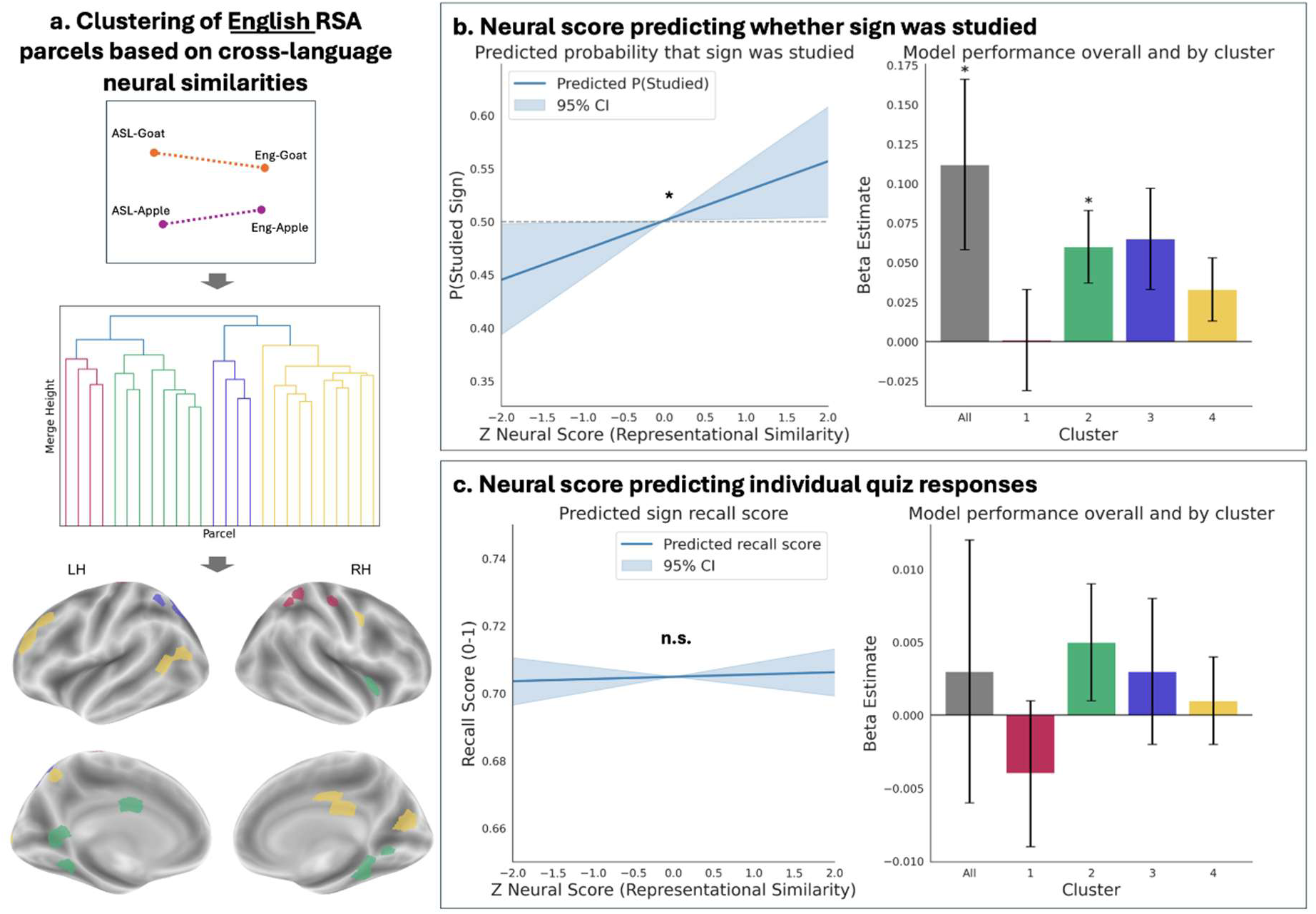
Cross-language similarity predicting individual differences in English RSA regions. Note. Linear mixed effect model results for English RSA regions. A. Diagram of hierarchical clustering procedure. Color-coded map of the parcels belonging to each of the four clusters is displayed on the semi-inflated FSAverage6 surface. B. Neural score predicting whether each sign belonged to the participant’s studied sign list. The change in predicted probability that the sign was studied over the range of neural scores is plotted on the left, while individual beta estimates of neural score for the entire model as well as split by cluster appear on the right. C. The same results are presented for neural score predicting individual recall quiz score. Bars represent standard error and are marked with asterisks by significance (p<0.05=*, p<0.01=**, p<0.001=***).

The overall model which included neural score as a fixed effect and participant, item, and cluster as random effects showed a modest relationship between neural score and whether the sign was studied (β=0.112, p=0.038). Overall and by-cluster results are summarized in Table 2. There was no significant prediction of individual recall score based on neural score in these regions.

**Table 2.** Linear Mixed Model Results for English RSA Regions.

Linear Mixed Model Results for English RSA Regions
| Model | $\beta$ | CI | Marginal $R^2$ | p-adj |
| --- | --- | --- | --- | --- |
| **Predicting studied sign** |  |  |  |  |
| All English Clusters | 0.112 | 0.01-0.22 | 0.001 | **0.038\*** |
| English Cluster 1 | 0.001 | -0.06-0.07 | >0.001 | 0.965 |
| English Cluster 2 | 0.060 | 0.02-0.10 | 0.001 | **0.036\*** |
| English Cluster 3 | 0.065 | 0.00-0.13 | 0.001 | 0.086 |
| English Cluster 4 | 0.033 | -0.01-0.07 | >0.001 | 0.134 |
| **Predicting recall score** |  |  |  |  |
| All English Clusters | 0.003 | -0.01-0.02 | >0.001 | 0.71 |
| English Cluster 1 | 0.001 | -0.06-0.07 | >0.001 | 0.725 |
| English Cluster 2 | 0.060 | 0.02-0.10 | 0.001 | 0.616 |
| English Cluster 3 | 0.065 | 0.00-0.13 | 0.001 | 0.725 |
| English Cluster 4 | 0.033 | -0.01-0.07 | >0.001 | 0.769 |

*Note. Normalized beta estimates, confidence intervals, marginal $R^2$ , and adjusted p-values are provided for the overall model as well as each within-cluster model for the regions selected with the English RSA. It should be noted that the prediction of studied from unstudied signs is a binomial (logistic) model, while recall score is predicted with a gaussian model, so effect sizes are not directly comparable between the two models.*

## Discussion

These results demonstrate that in hearing novice signers, greater similarity between the neural representation of a noun in ASL and its translational equivalent in English indicates not only familiarity with the sign, but individual ability to recall the sign’s meaning. This effect was seen in regions that also represented within-language conceptual information about the signs in ASL. However for regions which were correlated with those same semantic features in English, the prediction of which signs were studied was weaker and there was no evidence of recall prediction. Therefore, it is possible to detect shifts in neural representations of individual signs after as little as 45 minutes of training, and this effect may be driven primarily by areas recruited for semantic processing of signs (rather than speech).

Consistent with a previous study which utilized a nearly identical learning paradigm^56^, we find evidence for decoding one language in regions defined by semantic RSA correlation to the other language. This study takes an important step further by demonstrating that this shift, which happens remarkably early in the learning trajectory, is sensitive to individual learning as measured by a behavioral quiz. The counterbalanced design allows us to examine studied and unstudied signs within the same group of individuals with approximately 45 minutes of ASL training and show that studied signs are represented more similarly to their English counterparts than unstudied ones. We also show that individual item-level neural scores track behaviorally measured recall and can therefore be an informative measure of successful learning. Unlike previous investigations in this vein, our neural score measure does not rely on any *a priori* model of the stimuli (such as semantic distances in word2vec), instead directly comparing representational distance between translationally equivalent nouns.

The only other study to date which has utilized this type of analysis with sign language stimuli concluded that while there was evidence for shared representations of broad semantic information, fine-grained item information appeared to be language-specific^22^. However, unlike the bimodal bilinguals studied by Evans and colleagues, our participants were novices at the very earliest stage of the learning trajectory. Our results suggest that in this case, there are in fact shifts in individual item representations that indicate shared or coactivated semantic representations, and that the *degree* of this similarity is indicative of successful comprehension. This supports the account of language learning whereby novices utilize their established language to scaffold new learning^19,40^. Further research should probe further time points along the learning trajectory to explore whether neural similarity between sign and speech continues to increase with proficiency, and whether there is a point where late learners may develop a more “native-like” separation between the two languages. Some research suggests that univariate effects (for example, the recruitment of right parietal areas for spatial processing of sign^36,38^) might be sensitive to age of acquisition. These methods could provide a powerful way to test those dissociations.

Given the complex and multisensory nature of language processing, it is unsurprising that a distributed network of brain regions contribute to this effect. ASL Cluster 3, which includes three neural parcels (Schaefer 500-parcel atlas numbers 92, 155, and 247) in the left intraparietal sulcus, angular gyrus and precuneus showed the strongest evidence of both types of learning effects. Significant effects were also found in ASL Clusters 1 and 4, which encompass many areas in bilateral temporal lobes (including the left SMG, which was the region associated with the greatest cross-language overlap in the previous study^56^ and which is considered a central part of the perisylvian language network ^67,68^, including for sign^41^). Bilateral parcels in STS, another region often associated with sign processing, are also included in Cluster 4. Prior research suggests this area is involved in auditory processing of speech^69^ as well as biological motion processing^70^. For signed languages in particular, right STS may support processing of prosody and facial expression^71^. For each of these regions, our findings demonstrate that beyond shifts in univariate activation, multivariate patterns contain information that is predictive of fine-grained learning outcomes.

However, the strongest estimates resulted from the model which included all four clusters, indicating that the best prediction of behavioral scores came from considering the network as a whole rather than dividing it into component parts. The patterns in semantic representation that change as a result of learning are distributed across a wide set of neural regions and seem to be stronger in areas which support semantic processing of the new language (ASL) rather than the established language (English). Studies of novice spoken language learners have shown that the extent of neural similarity between their newly-learned and established languages increases with training and may correlate with proficiency^20,72,73^. Our findings go one step further, demonstrating that not just univariate overlap but subtle shifts in neural activity patterns contain information which correlates with individual ability to explicitly translate the semantic content of each sign to a familiar language.

These results show that among the novice learners who learned the most, multivariate pattern analysis techniques can detect shifts which differentiate not only signs they studied versus signs they did not, but individual likelihood of successful recall. The direct comparison between individual pairs of equivalent nouns suggests that unlike bimodal bilinguals, novices may represent fine-grained item distinctions in a way that is partially shared between languages, possibly due to coactivating their established language to aid translation^19,40^. Indeed, after such a short lesson paradigm, it is unlikely that participants had developed a nuanced and diverging representation of the ASL signs, but it is striking that this increase in representational similarity with learning occurs so rapidly and that they track individual behavioral scores. While the temporal features of this shift at the very early stages of learning remain understudied, these data provide evidence that it begins after less than an hour of instruction and might be supported more strongly by regions which show language-specific semantic processing for sign rather than those which support processing in the established language.

