## Supplemental Information for "Initial signs of learning: Decoding newly-learned vocabulary from neural patterns in novice sign language learners"

**Supplemental Figure 1.**

Group-level RSA results by language

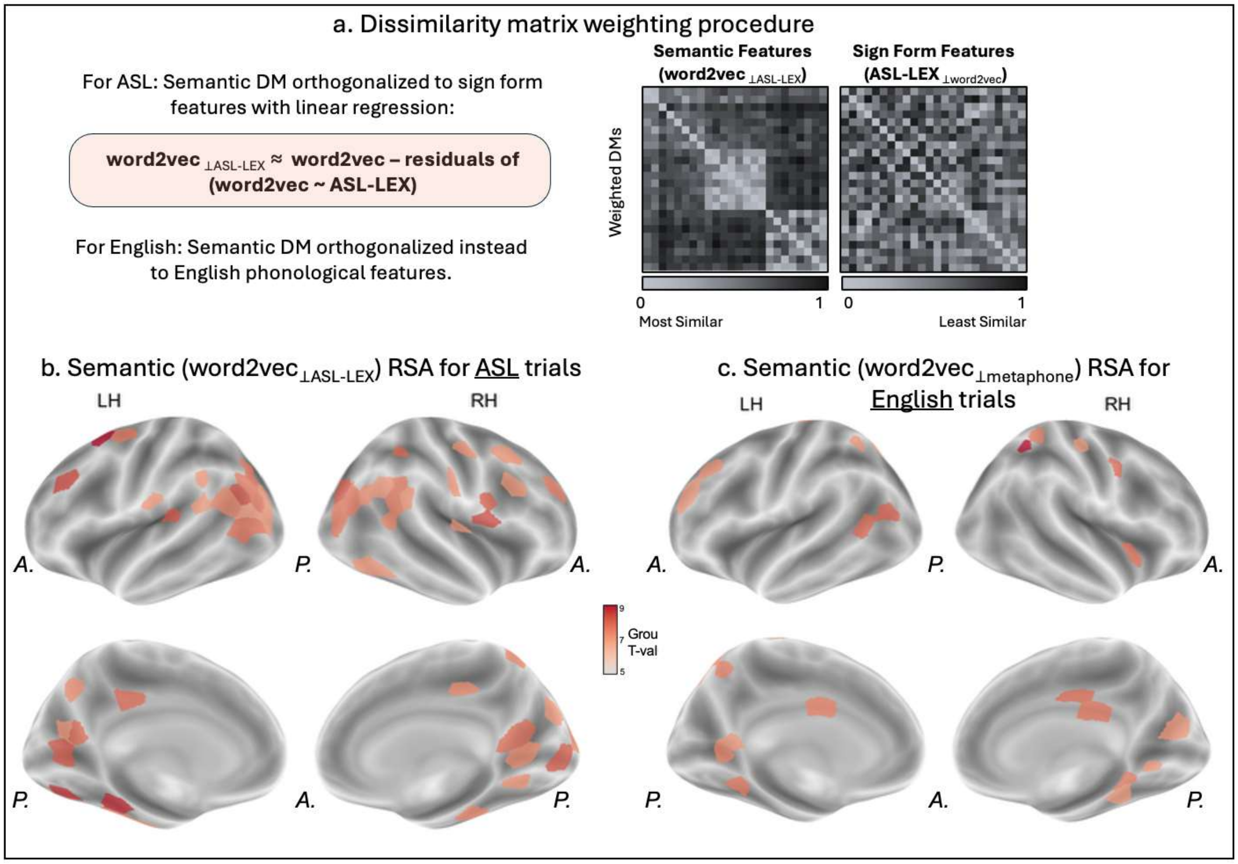


Note. Group RSA procedure. A. Dissimilarity matrix weighting procedure. The *a priori* models of semantic and phonological features were orthogonalized to one another using linear regression separately for each language. Thus, for ASL, semantic features derived from word2vec and sign form features derived from the ASL-LEX database were orthogonalized to each other before RSA. B. Semantic (word2vec) RSA results for ASL trials. Parcels with group-level corrected p-values < 0.05 are shown on the semi-inflated cortical surface (FSAverage6). The heatmap shows T-values of this group-level test. C. Semantic (word2vec) results for English trials, thresholded and plotted with the same procedure as the ASL trials. The parcels in B and C make up the binary masks used for subsequent individual differences analyses.

**Supplemental Table 1.**

Schaefer parcel labels for each cluster

| Cluster | Size (parcels) | Schaefer atlas labels (500-parcel version) |
|---|---|---|
| ASL Cluster 1 | 16 | 1, 2, 51, 54, 107, 108, 148, 200, 251, 260, 292, 302, 309, 346, 381, 383 |
| ASL Cluster 2 | 19 | 6, 27, 29, 32, 36, 90, 199, 239, 271, 274, 277, 278, 282, 336, 339, 367, 389, 491 |
| ASL Cluster 3 | 3 | 92, 155, 247 |
| ASL Cluster 4 | 16 | 156, 175, 201, 202, 203, 205, 245, 370, 384, 415, 437, 442, 453, 456, 457, 458 |
| English Cluster 1 | 4 | 85, 312, 318, 353 |
| English Cluster 2 | 8 | 7, 24, 182, 237, 253, 258, 265, 376 |
| English Cluster 3 | 4 | 96, 98, 101, 103 |
| English Cluster 4 | 10 | 25, 117, 128, 179, 199, 229, 280, 374, 387, 447 |

Note. Parcel count and identities for each hierarchical cluster defined by the English or ASL Group-level RSA. Parcel IDs refer to labels from the 500-parcel Schaefer atlas.

### fMRIprep Boilerplate Methods

This boilerplate text was automatically generated by fMRIprep with the express intention that users should copy and paste this text into their manuscripts unchanged. It is released under the CC0 license (https://creativecommons.org/publicdomain/zero/1.0/)

Preprocessing was performed using *fMRIprep* 21.0.1 (Esteban, Markiewicz, et al. 2018; Esteban, Blair, et al. 2018; RRID:SCR_016216), which is based on *Nipype* 1.6.1 (K. Gorgolewski et al. 2011; K. J. Gorgolewski et al. 2018; RRID:SCR_002502).

#### *Anatomical data preprocessing*

A total of 1 T1-weighted (t1w) images were found within the input BIDS dataset.The T1-weighted (t1w) image was corrected for intensity non-uniformity (INU) with n4biasfieldcorrection (Tustison et al. 2010), distributed with ants 2.3.3 (Avants et al. 2008, RRID:SCR_004757), and used as t1w-reference throughout the workflow. The t1w-reference was then skull-stripped with a *Nipype* implementation of the antsbrainextraction.sh workflow (from ants), using OASIS30ANTs as target template. Brain tissue segmentation of cerebrospinal fluid (CSF), white-matter (WM) and gray-matter (GM) was performed on the brain-extracted T1w using fast (FSL 6.0.5.1:57b01774, RRID:SCR_002823, Zhang, Brady, and Smith 2001). Brain surfaces were reconstructed using recon-all (freesurfer 6.0.1, RRID:SCR_001847, Dale, Fischl, and Sereno 1999), and the brain mask estimated previously was refined with a custom variation of the method to reconcile ants-derived and freesurfer-derived segmentations of the cortical gray-matter of Mindboggle (RRID:SCR_002438, Klein et al. 2017). Volume-based spatial normalization to one standard space (MNI152NLin2009cAsym) was performed through nonlinear registration with antsregistration (ants 2.3.3), using brain-extracted versions of both T1w reference and the T1w template. The following template was selected for spatial normalization: *ICBM 152 Nonlinear Asymmetrical template version 2009c* [Fonov et al. (2009), RRID:SCR_008796; templateflow ID: MNI152NLin2009cAsym].

***Functional data preprocessing***

For each of the 6 BOLD runs found per subject (across all tasks and sessions), the following preprocessing was performed. First, a reference volume and its skull-stripped version were generated using a custom methodology of *fMRIprep*. Head-motion parameters with respect to the BOLD reference (transformation matrices, and six corresponding rotation and translation parameters) are estimated before any spatiotemporal filtering using mcflirt (FSL 6.0.5.1:57b01774, Jenkinson et al. 2002). BOLD runs were slice-time corrected to 1.19s (0.5 of slice acquisition range 0s-2.37s) using 3dtshift from AFNI (Cox and Hyde 1997, RRID:SCR_005927). The BOLD time-series (including slice-timing correction when applied) were resampled onto their original, native space by applying the transforms to correct for head-motion. These resampled BOLD time-series will be referred to as *preprocessed BOLD in original space*, or just *preprocessed BOLD*. The BOLD reference was then co-registered to the T1w reference using bbregister (freesurfer) which implements boundary-based registration (Greve and Fischl 2009). Co-registration was configured with six degrees of freedom. Several confounding time-series were calculated based on the *preprocessed BOLD*: framewise displacement (FD), DVARS and three region-wise global signals. FD was computed using two formulations following Power (absolute sum of relative motions, Power et al. (2014)) and Jenkinson (relative root mean square displacement between affines, Jenkinson et al. (2002)). FD and DVARS are calculated for each functional run, both using their implementations in *Nipype* (following the definitions by Power et al. 2014). The three global signals are extracted within the CSF, the WM, and the whole-brain masks. Additionally, a set of physiological regressors were extracted to allow for component-based noise correction (*compcor*, Behzadi et al. 2007). Principal components are estimated after high-pass filtering the *preprocessed BOLD* time-series (using a discrete cosine filter with 128s cut-off) for the two *compcor* variants: temporal (tcompcor) and anatomical (acompcor). Tcompcor components are then calculated from the top 2% variable voxels within the brain mask. For acompcor, three probabilistic masks (CSF, WM and combined CSF+WM) are generated in anatomical space. The implementation differs from that of Behzadi et al. In that instead of eroding the masks by 2 pixels on BOLD space, the acompcor masks are subtracted a mask of pixels that likely contain a volume fraction of GM. This mask is obtained by dilating a GM mask extracted from the freesurfer's *aseg* segmentation, and it ensures components are not extracted from voxels containing a minimal fraction of GM. Finally, these masks are resampled into BOLD space and binarized by thresholding at 0.99 (as in the original implementation). Components are also calculated separately within the WM and CSF masks. For each compcor decomposition, the *k* components with the largest singular values are retained, such that the retained components' time series are sufficient to explain 50 percent of variance across the nuisance mask (CSF, WM, combined, or temporal). The remaining components are dropped from consideration. The head-motion estimates calculated in the correction step were also placed within the corresponding confounds file. The confound time series derived from head motion estimates and global signals were expanded with the inclusion of temporal derivatives and quadratic terms for each (Satterthwaite et al. 2013). Frames that exceeded a threshold of 0.5 mm FD or 1.5 standardised DVARS were annotated as motion outliers. The BOLD time-series were resampled into standard space, generating a *preprocessed BOLD run in MNI152NLin2009cAsym space*. First, a reference volume and its skull-stripped version were generated using a custom methodology of *fMRIprep*. The BOLD time-series were resampled onto the following surfaces (freesurfer reconstruction nomenclature): *fsnative*. All resamplings can be performed with *a single interpolation step* by composing all the pertinent transformations (i.e. Head-motion transform matrices, susceptibility distortion correction when available, and co-registrations to anatomical and output spaces). Gridded (volumetric) resamplings were performed using antsapplytransforms (ants), configured with Lanczos interpolation to minimize the smoothing effects of other kernels (Lanczos 1964). Non-gridded (surface) resamplings were performed using mri_vol2surf (freesurfer).

Many internal operations of *fMRIprep* use *Nilearn* 0.8.1 (Abraham et al. 2014, RRID:SCR_001362), mostly within the functional processing workflow. For more details of the pipeline, see the section corresponding to workflows in *fMRIprep*'s documentation (https://fMRIprep.readthedocs.io/en/latest/workflows.html).

**fMRIprep References**


Abraham, Alexandre, Fabian Pedregosa, Michael Eickenberg, Philippe Gervais, Andreas Mueller, Jean Kossaifi, Alexandre Gramfort, Bertrand Thirion, and Gael Varoquaux. 2014. "Machine Learning for Neuroimaging with Scikit-Learn." *Frontiers in Neuroinformatics* 8. Https://doi.org/10.3389/fninf.2014.00014.

Avants, B. B., C. L. Epstein, M. Grossman, and J. C. Gee. 2008. “Symmetric Diffeomorphic Image Registration with Cross-Correlation: Evaluating Automated Labeling of Elderly and Neurodegenerative Brain.” *Medical Image Analysis* 12 (1): 26–41. Https://doi.org/10.1016/j.media.2007.06.004.

Behzadi, Yashar, Khaled Restom, Joy Liau, and Thomas T. Liu. 2007. “A Component Based Noise Correction Method (compcor) for BOLD and Perfusion Based fMRI.” *Neuroimage* 37 (1): 90–101. Https://doi.org/10.1016/j.neuroimage.2007.04.042.

Ciric, R., William H. Thompson, R. Lorenz, M. Goncalves, E. Macnicol, C. J. Markiewicz, Y. O. Halchenko, et al. 2022. “templateflow: FAIR-Sharing of Multi-Scale, Multi-Species Brain Models.” *Nature Methods* 19: 1568–71. Https://doi.org/10.1038/s41592-022-01681-2.

Esteban, Oscar, Ross Blair, Christopher J. Markiewicz, Shoshana L. Berleant, Craig Moodie, Feilong Ma, Ayse Ilkay Isik, et al. 2018. “fMRIprep 24.1.1.” *Software*. Https://doi.org/10.5281/zenodo.852659.

Esteban, Oscar, Christopher Markiewicz, Ross W Blair, Craig Moodie, Ayse Ilkay Isik, Asier Erramuzpe Aliaga, James Kent, et al. 2019. “fMRIprep: A Robust Preprocessing Pipeline for Functional MRI.” *Nature Methods* 16: 111–16. Https://doi.org/10.1038/s41592-018-0235-4.

Fonov, VS, AC Evans, RC McKinstry, CR Almli, and DL Collins. 2009. “Unbiased Nonlinear Average Age-Appropriate Brain Templates from Birth to Adulthood.” *Neuroimage* 47, Supplement 1: S102. Https://doi.org/10.1016/S1053-8119(09)70884-5.

Gorgolewski, K., C. D. Burns, C. Madison, D. Clark, Y. O. Halchenko, M. L. Waskom, and S. Ghosh. 2011. “Nipype: A Flexible, Lightweight and Extensible Neuroimaging Data Processing Framework in Python.” *Frontiers in Neuroinformatics* 5: 13. Https://doi.org/10.3389/fninf.2011.00013.

Gorgolewski, Krzysztof J., Oscar Esteban, Christopher J. Markiewicz, Erik Ziegler, David Gage Ellis, Michael Philipp Notter, Dorota Jarecka, et al. 2018. "Nipype." *Software*. Https://doi.org/10.5281/zenodo.596855.

Greve, Douglas N, and Bruce Fischl. 2009. "Accurate and Robust Brain Image Alignment Using Boundary-Based Registration." *Neuroimage* 48 (1): 63–72. Https://doi.org/10.1016/j.neuroimage.2009.06.060.

Jenkinson, Mark, Peter Bannister, Michael Brady, and Stephen Smith. 2002. "Improved Optimization for the Robust and Accurate Linear Registration and Motion Correction of Brain Images." *Neuroimage* 17 (2): 825–41. Https://doi.org/10.1006/nimg.2002.1132.

Jenkinson, Mark, and Stephen Smith. 2001. "A Global Optimisation Method for Robust Affine Registration of Brain Images." *Medical Image Analysis* 5 (2): 143–56. Https://doi.org/10.1016/S1361-8415(01)00036-6.

Patriat, Rémi, Richard C. Reynolds, and Rasmus M. Birn. 2017. "An Improved Model of Motion-Related Signal Changes in fMRI." *Neuroimage* 144, Part A (January): 74–82. Https://doi.org/10.1016/j.neuroimage.2016.08.051.

Power, Jonathan D., Anish Mitra, Timothy O. Laumann, Abraham Z. Snyder, Bradley L. Schlaggar, and Steven E. Petersen. 2014. "Methods to Detect, Characterize, and Remove Motion Artifact in Resting State fMRI." *Neuroimage* 84 (Supplement C): 320–41. Https://doi.org/10.1016/j.neuroimage.2013.08.048.

Reuter, Martin, Herminia Diana Rosas, and Bruce Fischl. 2010. "Highly Accurate Inverse Consistent Registration: A Robust Approach." *Neuroimage* 53 (4): 1181–96. Https://doi.org/10.1016/j.neuroimage.2010.07.020.

Satterthwaite, Theodore D., Mark A. Elliott, Raphael T. Gerraty, Kosha Ruparel, James Loughead, Monica E. Calkins, Simon B. Eickhoff, et al. 2013. "An improved framework for confound regression and filtering for control of motion artifact in the preprocessing of resting-state functional connectivity data." *Neuroimage* 64 (1): 240–56. Https://doi.org/10.1016/j.neuroimage.2012.08.052.

Tustison, N. J., B. B. Avants, P. A. Cook, Y. Zheng, A. Egan, P. A. Yushkevich, and J. C. Gee. 2010. "N4ITK: Improved N3 Bias Correction." *IEEE Transactions on Medical Imaging* 29 (6): 1310–20. Https://doi.org/10.1109/TMI.2010.2046908.

Zhang, Y., M. Brady, and S. Smith. 2001. "Segmentation of Brain MR Images Through a Hidden Markov Random Field Model and the Expectation-Maximization Algorithm." *IEEE Transactions on Medical Imaging* 20 (1): 45–57. Https://doi.org/10.1109/42.906424.
